# Fluctuating selection facilitates the discovery of broadly effective but difficult to reach adaptive outcomes in yeast

**DOI:** 10.1101/2023.06.17.545402

**Authors:** Vincent J. Fasanello, Ping Liu, Justin C. Fay, Carlos A. Botero

## Abstract

Evolutionary compromises are thought to be common under fluctuating selection because the mutations that best enable adaptation to one environmental context can often be detrimental in others. Yet, prior experimental work has shown that generalists can sometimes perform as well as specialists in their own environments. Here we use a highly replicated evolutionary experiment (N = 448 asexual lineages of the brewer’s yeast) to show that even though fluctuation between two environmental conditions often induces evolutionary compromises (at least early on), it can also help reveal difficult to reach adaptive outcomes that ultimately improve performance in both environments. Specifically, we begin by showing that yeast adaptation to chemical stress can involve fitness tradeoffs with stress-free environments and that, accordingly, lineages that are repeatedly exposed to occasional stress tend to respond by trading performance for breadth of adaptation. We then show that on rare occasions, fluctuating selection leads to the evolution of no-cost generalists that can even outcompete constant selection specialists in their own environments. We propose that the discovery of these broader and more effective adaptive outcomes under fluctuating selection could be partially facilitated by changes in the adaptive landscape that result from having to deal with fitness tradeoffs across different environmental conditions. Overall, our findings indicate that reconciling the short- and long-term evolutionary consequences of fluctuating selection could significantly improve our understanding of the evolution of specialization and generalism.

## INTRODUCTION

The ability to persist under a wide range of environmental conditions is presumably advantageous because it could dramatically multiply a lineage’s opportunities on Earth. What then prevents the widespread evolution of generalists? The most common explanation for the relative rarity of generalists is that niche breadth tends to trade off with peak fitness, resulting in a cost to generalization [1]. However, empirical studies have often failed to detect such trade-offs [2–4], have found it difficult to distinguish them from conditional neutrality [4], or have failed to detect the actual performance dimensions that trade off against each other [1]. Alternatively, the evolution of generalism could also be constrained by comparatively faster rates of evolution toward specialization [5,6].

Experimental evolution has long been used to investigate the evolutionary process, particularly as it relates to exposure to heterogeneous conditions. While these experiments generally indicate that constant environments tend to promote the evolution of specialists and variable ones tend to promote generalization, there have also been reports of populations that respond to constant but not variable treatments [4,7]. Additionally, the grain of environmental variation, the type of variation (spatial or temporal), and the complexity and predictability of the environment, have all been shown to impact rates of adaptation and adaptive outcomes [7–17]. Thus, meaningful experimental investigations of the potential costs of generalism will require a proper standardization of ecological contexts. Comparing the evolution of identical microbial lineages under constant and fluctuating selection is a promising way to achieve such standardization.

In this study we examine yeast evolution in constant and temporally variable environments. Specifically, we evolve 448 genetically barcoded *Saccharomyces cerevisae* lineages in 14 treatments that vary in the concentration and dynamics of two chemical stresses: salt and copper. We find that adaptation to constant environments results in fitness trade-offs for salt but not copper treatments and that most populations that evolve in fluctuating treatments with either salt or copper trade, as expected, performance for breadth of adaptation. Additionally, we find that such trade-offs are not universal, as some fluctuating selection populations in our experiment were able to improve performance under every condition in their treatment and even evolved higher fitness specialists in their home environments.

## RESULTS

### Fitness gains in 500-generations of experimental evolution

We expected yeast lineages to adapt to the chemical stresses to which they were exposed. Accordingly, lineages that experienced less chemical stress during experimental evolution tended to exhibit greater fitness gains in the 0% stress environment (Fig 1A and 1B), whereas those that experienced higher stress concentrations during evolution tended to exhibit greater fitness gains in media with 80% chemical stress (Fig 1C and 1D).

**Figure 1:**
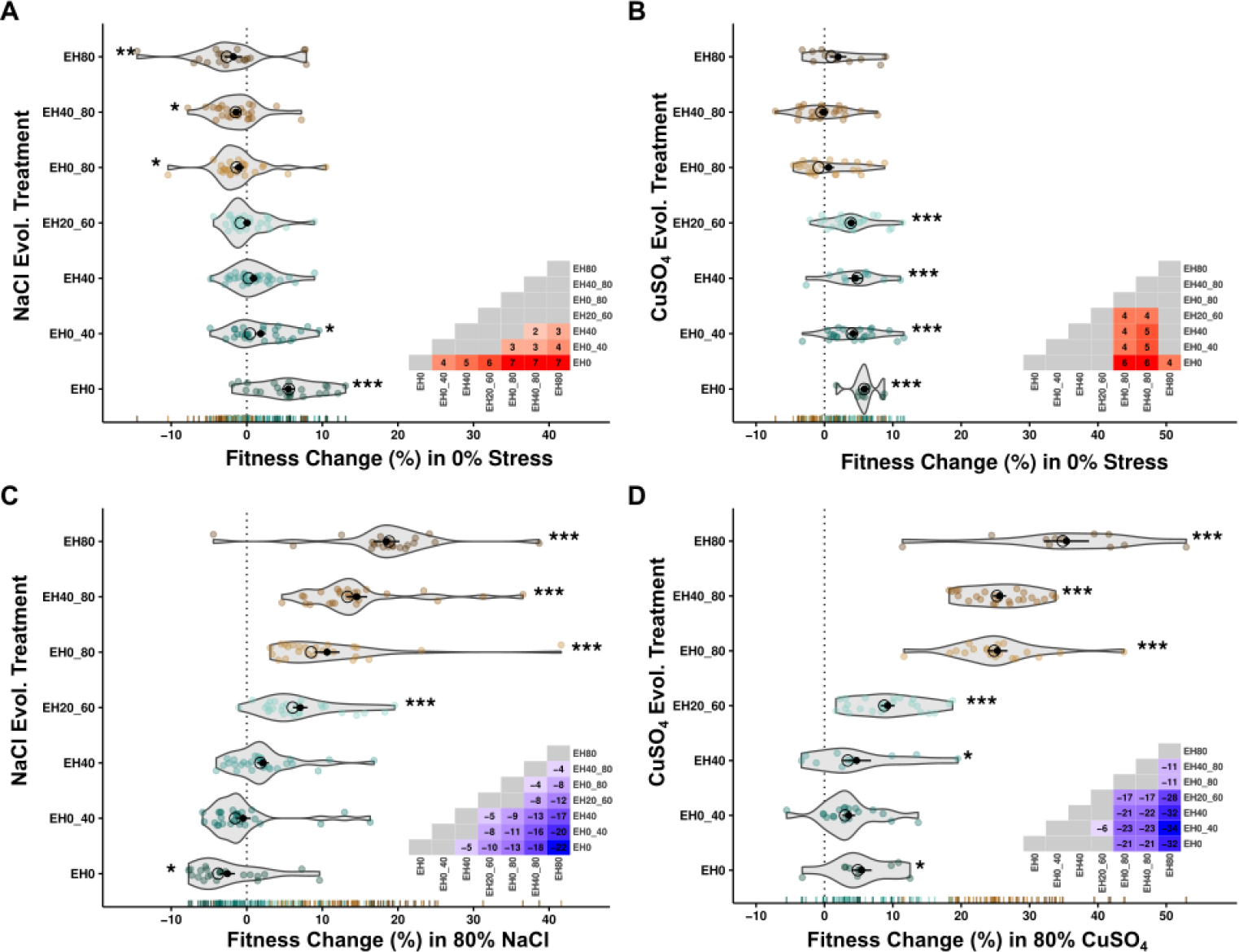
Fitness change in in 0%, 80% chemical stress after 500-generations experimental evolution. (A) Fitness change in 0% stress for lineages from the salt dataset. (B) Fitness change in 0% stress for the copper dataset. (C) Fitness change in 80% salt stress. (D) Fitness change in 80% copper stress. Lower-triangle insets illustrate fitness differences among treatments; significant associations have beta-values, non-significant associations in grey. Asterisks denote treatment differences from 0 (no fitness change). Black open circles are treatment median fitness. Black Closed circles are treatment mean fitness with standard error bars depicted.

Adaptation to a particular environmental condition is likely to impact fitness in other environmental conditions. These effects can be complementary, neutral, or antagonistic. We observed all three cases in our data. In the salt dataset, adaptation to 0% chemical stress was associated with negative fitness change in 80% stress (Fig 1A and 1C -EH0) and adaptation to 80% stress was associated with negative fitness change in the 0% stress environment (Fig 1A and 1C – EH80, EH40_80, EH0_80). In the copper dataset, adaptation to 0% chemical stress frequently resulted in a fitness increase in 80% stress (Fig 1B and 1D -EH0), while adaptation to 80% stress had neither positive nor negative effects on fitness in the 0% chemical stress environment (Fig 1A and 1C – EH80, EH40_80, EH0_80).

### Genetic correlation in fitness

Selection under uniform environmental conditions favors individuals whose fitness is highest in that environment and should therefore result in the evolution of specialists [7]. Consequently, cross-environment genetic correlation in fitness should evolve to become negative if adaptive mutations perform better in their evolutionary environment than in other environments. Positive genetic correlations in fitness arise when mutations perform equally or better in an alternative environment than lineages that actually evolved in such environment [7]. We tested for cross-environment genetic correlation in fitness by comparing the fitness of strains evolved under different experimental treatments in media with 0%, 40% and 80% stress.

As in Kassen [7], we identified cross-environment genetic correlations in fitness as the slope of the line connecting the fitness of the constant treatments for that pair of environmental conditions (Fig 2). Despite noteworthy differences in patterns of adaptation in the salt and copper datasets (Fig 1), patterns of cross-environment genetic correlation were qualitatively similar under both chemical stressors. Specifically, we observed negative cross-environment genetic correlations in fitness between 0% and 40% chemical stress (Fig 2A) and between 0% and 80% chemical stress for both stress treatments (Fig 2B). The slope of this relationship become more negative at increasing environmental dissimilarities (i.e., compare Fig 2A and 2B). In contrast, we found a positive cross-environment genetic correlation in fitness between the 40% stress and 80% stress environments for both salt and copper treatments (Fig 2C).

**Figure 2:**
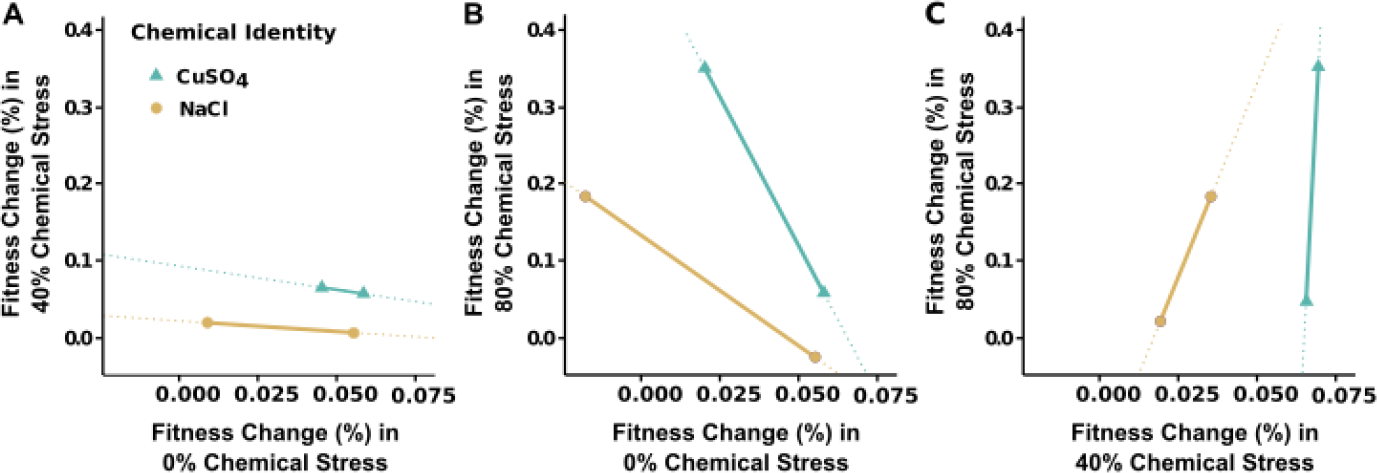
Cross-environment genetic correlation in fitness. Fitness change for each treatment is shown by the fitness change in the lower stress environment on the x-axis and fitness change in the higher stress environment on the y-axis. Negative slopes indicate the presence of a negative cross-environment genetic correlation in fitness change. Positive slopes indicate the presence of a positive correlation. (A) Blue. EH40, EH0 in 0%, 40% stress; (B) Purple. EH0, EH80 in 0%, 80% stress; (C) Red. EH40, EH80 in 40%, 80% stress. Salt dataset depicted with circle endcaps; copper dataset depicted with triangular endcaps. Dotted lines included for visual comparison of slopes. Axes have the same scaling in A, B, C to allow comparison of cross-environment genetic correlation in fitness between pairs of environments.

### Costs of adaptation in constant conditions

Negative cross-environment genetic correlation in fitness can evolve due to fitness trade-offs, in which adaptation to one environment has a fitness cost in others. Alternatively, negative cross-environment genetic correlation in fitness can arise in the absence of strict costs if direct responses to selection are larger than correlated responses in other environments, i.e. if each strain is more closely adapted to its home environment than to other conditions [18].

To evaluate the prevalence of costs of adaptation in our data, we assessed whether strains evolved under constant conditions exhibited fitness gains in their respective home environments and losses in other environments. When examining extremes at the treatment level—i.e., treatments EH0 and EH80 under 0% and 80% chemical stress—we observed costs in the salt dataset but not in the copper dataset (Fig 2). Results for individual lineages replicated these findings, indicating that costs were relatively common with salt (Fig 3A and 3C; Table S1), but rare with copper (Fig 3D and 3F; Table S1). Our results for the EH0 and EH80 treatments in 40% stress were intermediate (Fig 3A, 3C, 3D, 3F; Table S1). In fact, the majority of lineages from the EH40 treatments did not exhibit evidence of costs at 0% or 80% stress, regardless of the chemical stressor (Fig 3B, 3E; Table S1).

**Figure 3:**
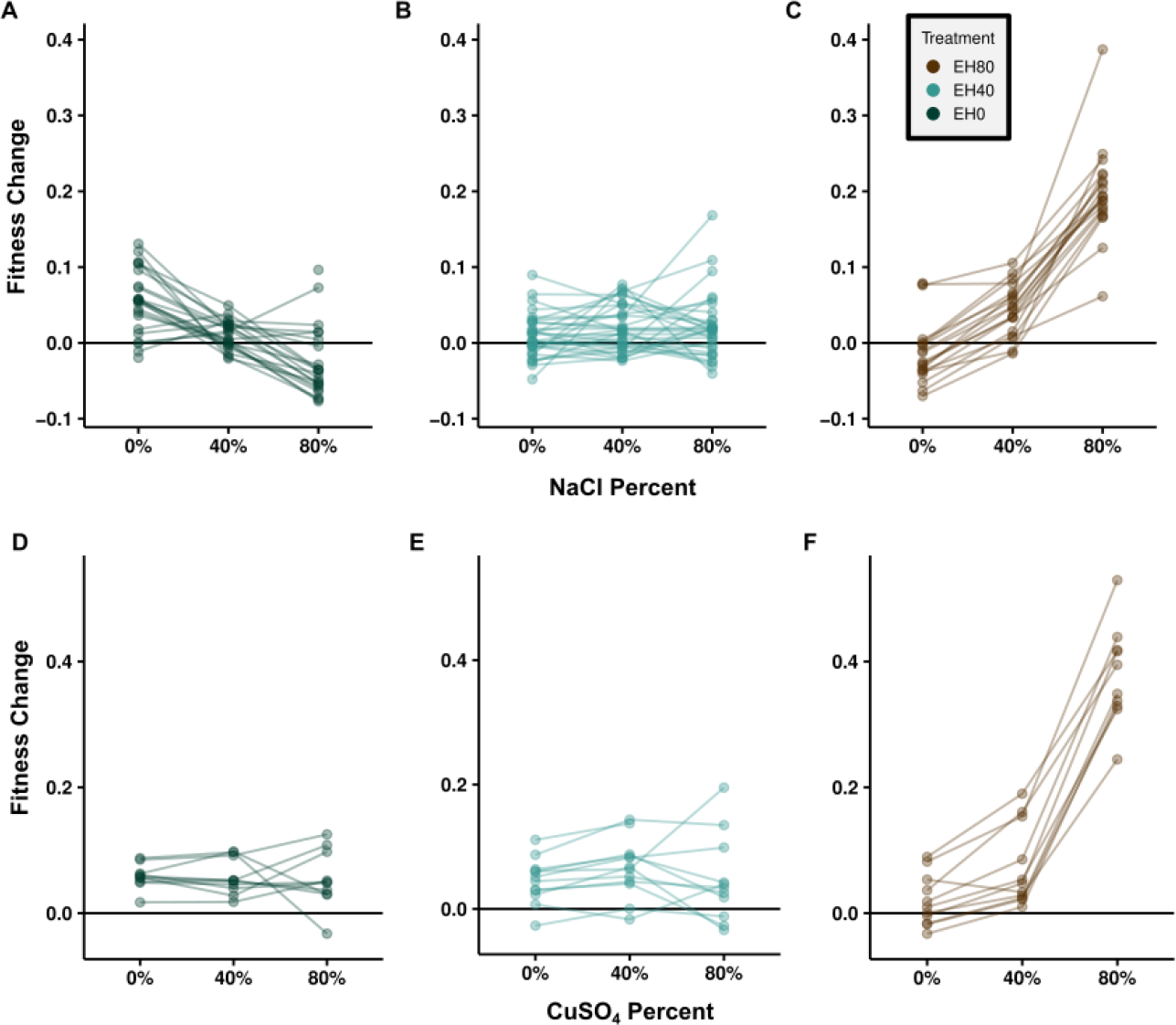
Fitness relationships for individual barcoded lineages. Fitness change in CM plus 0%, 40%, and 80% chemical stress for treatments exposed to constant chemical stress at 0% (EH0), 40% (EH40), and 80% (EH80) the ancestral lethal limit. Each line depicts the fitness change in three assay environments for each barcoded yeast population. Top row shows data from our salt dataset: (A) EH0, (B) EH40, (C) EH80. Bottom row shows data from our copper dataset: (D) EH0, (E) EH40, (F) EH80. One barcoded strain with low fitness and missing data removed from (A) and one strain with low fitness removed from (C) for visualization purposes only.

### Costs of adaptation in fluctuating conditions

It is also possible that the cost of generalization lies not in the existence of evolutionary tradeoffs between environments but rather on a slower rate of adaptation [7]. Under that scenario, generalists could theoretically improve fitness in both environments if given enough time. To evaluate whether trade-offs exist in this system, we assayed the fitness of lines selected in constant (EH0, EH80) and fluctuating conditions (EH0_40, EH20_60, EH0_80, EH40_80) in environments with 0% and 80% stress. Under this model, the absence of a trade-off between breadth of adaptation and performance could be inferred from the observation that populations evolved under fluctuating selection lie on the line connecting the performance of populations evolved under the EH0 and EH80 treatments. Alternatively, if breadth of adaptation increases at the expense of performance, then lineages evolved under fluctuating selection should fall below this line.

#### Costs are universal at the treatment level

All treatments with evolutionary histories in fluctuating environments (EH0_40, EH20_60, EH0_80, EH40_80) generally traded performance for breadth of adaptation in the copper data—i.e., the mean values for all treatments fell significantly below of the no-cost line at a detection limit of 0.634% fitness change (Fig 4B, 4D; Table S2, Methods). Although treatments EH0_40, EH20_60, and EH0_80 also exhibited this trade-off in the salt dataset, results were non-significant for the EH40_80 treatment (Fig 4A, 4C; Table S3). The magnitude of realized cost (distance below the no-cost line) was negatively associated with the amount of chemical stress experienced during evolution in the salt dataset (Fig 4A; Table S3). The opposite pattern exists in the copper data, where cost was positively associated with the amount of chemical stress experienced (Fig 4B; Table S2). Costs were also observed for the EH0_40 treatment relative to the constant EH0 and EH40 treatments for both chemicals (Fig S3).

**Figure 4:**
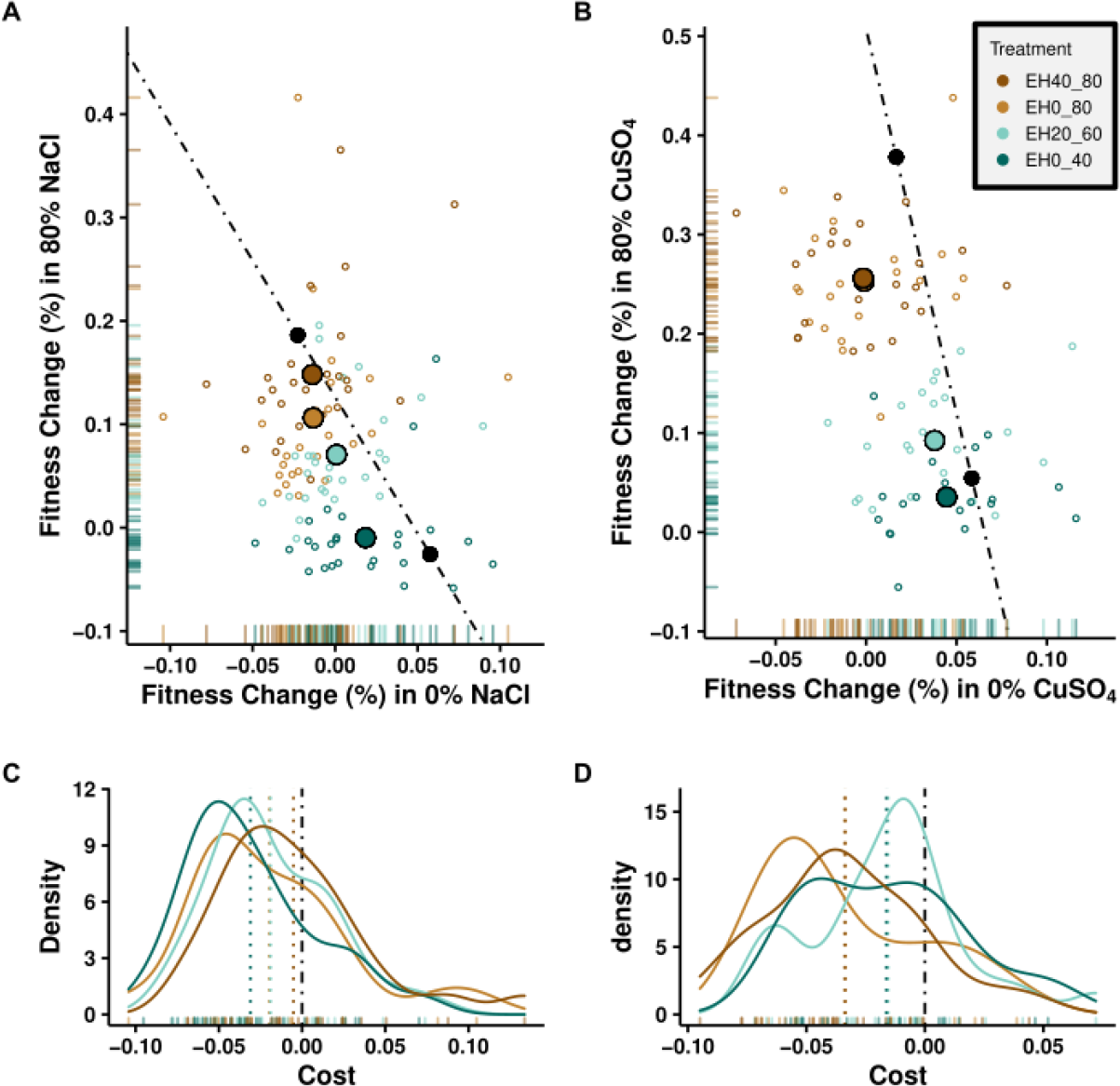
Lineages with evolutionary histories in fluctuating chemical stress environments do not always trade depth for breadth of adaptation. (A, B) Black circles are the treatment mean fitness for the constant chemical stress lineages EH0 (bottom, right) and EH80 (top, left). Orthogonal distance below- and to the left of the dot-dash line connecting the EH0 and EH80 indicates a fitness cost in the form of a trade-off of depth for breadth of adaptation. Lineages on the line pay no cost of generalization. Lineages orthogonally above- or to the right of the dot-dash line enjoy additional fitness benefits in the form of depth and breadth of adaptation. (A,B) Cost of generalism for all fluctuating chemical stress treatments (EH0_40, EH20_60, EH0_80, EH40_80) relative to the constant 0% (EH0) and 80% chemical stress (EH80) treatments. (C,D) Corresponds to (A,B) data; density plots depict cost, distance above (+), below (-) the dot-dash line. (A,C) data for the salt dataset; (B,D) copper dataset.

#### A diversity of outcomes exists within treatments

Coarse, treatment-level, results may suggest the general existence of a trade-off between breadth of adaption and performance but fail to capture the full range of adaptive outcomes that are observable within treatments. For example, examination of fitness changes for individual lineages revealed a rich diversity of phenotypes within each treatment and uncovered broad overlap in fitness phenotypes among treatments (Figure 4, small open circles). For example, in the salt data, 58/105 (55%) of populations across the EH0_40, EH20_60, EH0_80, and EH40_80 treatments exhibited significant costs at a detection limit of 2.163% fitness change (Methods). In the copper dataset, the number was 46/88 (82%). In both datasets, we also found a subset of replicates that exhibit no evidence of a trade-off given that their performance was not significantly different from the no-cost line connecting the EH0 and EH80 treatments (i.e., 30/105 or 29% of replicates in the salt dataset and 34/88 or 39% of replicates in the copper dataset). Furthermore, we also found that every treatment contained at least some lineages that fell significantly above the no cost line, indicating that some lineages may have developed adaptations that allowed them to generally increase performance across the range of fluctuating conditions (i.e., 17/105 or 16% of replicates in the salt dataset, and 8/88 or 9% of replicates in the copper dataset, Table S4).

## DISCUSSION

Generalists are often depicted as “jacks of all trades [and] masters of none” [19] because exploiting a range of environmental conditions can often involve compromise. We examined the potential for antagonistic pleiotropy in yeast by exposing a variety of lineages to constant selection from either salt or copper and found that stress-adapted populations exhibit clear fitness trade-offs only in the salt context. Although the negative fitness correlations reported in Fig 2 are often interpreted as evidence of antagonistic pleiotropy, we note that in some instances they could also arise from (cost-free) conditional mutations that are beneficial in one environment but neutral [4] or less effective [18] in others. Thus, although uncertainty in the actual mechanism remains, our findings confirm that specialization to one context can impact performance in others.

The notion that effective exploitation of a given environmental condition can compromise performance in other contexts is further supported by our finding that lineages exposed to fluctuating selection evolved to be less fit in complete media and under chemical stress than lineages exclusively evolved under either of those conditions (Fig 4). The fact that such compromises are evident even under copper stress (a treatment that showed no evidence of fitness tradeoffs), demonstrates that the costs of generalism in yeast are not exclusively related to antagonistic pleiotropy (or conditional performance). A likely alternative is that fluctuating selection lineages were unable to match the performance of constant selection lineages in our experiment because they exhibit slower rates of evolution [5–7] and had simply not yet fully adapted. Regardless of the mechanism, our findings demonstrate that initial adaptation to fluctuating selection in yeast involves adaptive compromises that generally reduce performance across the range of environmental conditions that these organisms experience (a likely version of bet-hedging [20]).

We emphasize that our findings relate to the initial stages of adaptation to fluctuating selection because our experiment was relatively short (500 generations). As populations evolve closer to their optimum, antagonistic pleiotropy could, for example, become more important [21,22] even in the copper context. Additionally, the sequential evolution of compensatory changes or epistasis could eventually enable cost-free generalism [2,4]. As a matter of fact, the large number of replicates in this study allowed us to demonstrate that such no-cost generalist mutants do evolve in this system, albeit on rare occasions. The rarity of this outcome could be related to the fact that mutations that confer advantages in both environments are either rare themselves or have smaller individual effects [23]. In either case, the competitive superiority of these mutants in both chemical contexts of our experiment suggests that no-cost generalists may be relatively common in natural populations of the brewer’s yeast, where both the genetic diversity and the timeframe of evolution are much larger than in our experimental setup.

In conclusion, our findings demonstrate that initial adaptation to fluctuating selection can be challenging because many mutations that confer advantages in one environmental context can be detrimental in others. Nevertheless, our experiment also demonstrates that exposure to oscillating conditions can sometimes result in the evolution of generalists that can even outcompete specialists in their respective home environments. This latter finding is particularly significant because it suggests that fluctuating selection can facilitate the discovery of broadly adaptive traits that are effectively inaccessible under constant selection. One potential way in which these discoveries may be facilitated is by modifying the adaptive landscape in a way that more readily reveals an evolutionary path to those more favorable trait combinations. For example, geometric mean fitness under fluctuating selection could smooth or perhaps even remove fitness peaks or valleys that in constant selection backgrounds drive populations away from these broadly favorable trait combinations. Thus, our findings suggest overall that a better reconciliation of the short- and long-term evolutionary consequences of fluctuating selection may significantly improve our understanding the evolution of specialization and generalism.

## METHODS

### Strains, media and culture methods

Barcoded yeast strains were constructed using two isogenic haploid derivatives of a strain collected from an oak tree in Pennsylvania (YPS163) [24]: YJF153 (MATa, *HO*::dsdAMX4) and YJF154 (MATalpha, *HO*::dsdAMX4) [25]. 113 diploid strains (Table S5) were constructed such that each contained a unique 20bp barcode-sequence flanking KAN, inserted in the HO locus [26]. A single barcoded strain (d1H10) was arbitrarily selected from this set to serve as a reference; the remaining 112 barcoded strains were subject to 50 days of experimental evolution (See *Experimental design*, below).

Yeast were cultured in complete medium (CM; 20 g/l dextrose, 1.7 g/l yeast nitrogen base without amino acid and ammonium sulfate, 5.0 g/l ammonium sulfate, 1.3 g/l dropout mix complete without yeast nitrogen base) with or without additional chemical stress in 96-deep well plates (2.2-ml poly-propylene plates, square well, v-conical bottoms; Abgene AB-0932) covered with rayon acrylate breathable membranes (Thermo Scientific, 1257605). Growth plates were incubated at 30°C for 24 hours inside an incubator (VWR, Forced Air Incubator, basic, 120v, 7 cu. ft.) with agitation using a horizontal electromagnetic microplate shaker (Union Scientific LLC, 9779-TC). Saturated (stationary phase) 24-hour culture was diluted (1:1000) into fresh medium at the same time each day to initialize the next round of growth.

### Experimental design

The experimental design included a 50-day experimental evolution with subsequent fitness quantification of ancestral (Day-0) and evolved (Day-50) yeast via competition-based fitness assay.

#### Experimental Evolution

112 barcoded yeast strains were divided evenly among seven treatments variable for chemical stress concentration and temporal dynamics. Constant chemical stress treatments were evolved for 50 days in Complete Medium (CM) plus chemical stress at 0% (EH0, read as: Evolutionary History 0%), 40% (EH40), or 80% (EH80) of the lethal limit for unevolved yeast strains in our library. The ancestral lethal limit for salt was 20 g/l NaCl and for copper was 8 um CuSO_4_. For these constant environments chemical stress concentration did not change from transfer-to-transfer. Fluctuating treatments were evolved for 50 days in chemical stress that alternated daily between two concentrations: 0%-40% (EH0_40), 20%-60% (EH20_60), 40%-80% (EH40_80), or 0%-80% (EH0_80) of the ancestral limit. This design was copied to create four microplates which were evolved in parallel for 50 days: two were exposed to salt stress and two were exposed to copper stress. Stress concentrations were selected such that they were comparable between chemical stressors and such that the 80% stress treatment reduced growth but did not result in extinction (from transfer to transfer) for an average ancestral strain. Samples were collected from the initial mixtures (starting material for plate copies, Day-0) and from the final overnight cultures (on Day-50). These samples served as the starting material for the Day-0 and Day-50 fitness assays, respectively.

#### Fitness Assays

Sequencing based competition assays, hereafter fitness assays, were subsequently conducted on Day-0 (ancestral) and Day-50 (evolved) yeast to assess fitness relative to the reference strain. Yeast lines from Day-0 and Day-50 of the experimental evolution were revived from stocks and mixed, separately, in equal proportions to create five pools. A single pool was created from each [evolutionary microplate] X [day] for a total of 1 Day-0 sample (the template for the evolution) and 4 Day-50 samples (2 salt evolved and 2 evolved in copper). The reference strain was then spiked into each pool at a high proportion (∼70%). Pools were diluted into fresh medium and cultured for two rounds of growth to allow competition to occur. Fitness assays were conducted in CM with and without additional chemical stress. Day-0 and Day-50 Fitness assays were run in quadruplicate and initial measures (barcode starting proportions) for each were taken in quintuplicate. Samples were collected from the initial mixtures (fitness assay starting material) and from the final cultures (ca. 20 generations later). From these data, the fitness of each barcoded line prior to evolution (Day-0 assays) and after evolution (Day-50 assays) was quantified and the resulting values were used to assess change in fitness for each line in each environment relative to the reference (see *Fitness Calculations*, below).

### Library construction and sequencing

DNA was isolated using a ZR Fungal/Bacterial DNA Kit (Zymo Research D6005) in individual 2.0 mL screw-cap tubes following the manufacturer’s instructions. Physical cell disruption by bead-beating was conducted in a mixer mill (Retsch, MM 300) at 30 Hz (1800 min^-1^) for ten minutes (1-minute on, 1-minute off, times ten cycles). MoBY barcodes were then amplified with forward/reverse Ion Torrent adapters containing a 9-12 bp index for multiplex sequencing (Table S6). PCR products for library construction were generated using 25 cycles and were subsequently quantified with a Qubit 3.0 Flourometer (ThermoFisher Scientific, Q33216) using the high sensitivity assay kit (ThermoFisher Scientific, Q32851). Products were combined at equimolar concentrations and purified using a Zymo DNA Clean and Concentrator kit (Zymo Research D4014) to create a single multiplexed library for sequencing. Additional control samples were included in the library to track barcode cross-contamination as well as any contamination that may have occurred during sample processing. An aliquot of the library was sequenced using an Ion Torrent sequencer (Ion Proton System, Ion Torrent) at the Genomics Core Facility at Saint Louis University with a customized parameter to assess polyclonality after 31bp (the start of the forward Ion Torrent adapter index sequence). A second aliquot was sequenced to augment read depth following preliminary assessment of data quality.

### Sequence data processing and calculations

#### Sequence dataset

Sequence data in FASTQ format were parsed and demultiplexed using custom scripts in R. 96,807,316 reads were retained for analysis that perfectly matched a forward adapter index (9-12 bp), a reverse adapter index (9-12 bp), and a MoBY genetic barcode (20 bp) included in the full experimental design. Of these, 52,853,350 (54.6%) mapped to non-reference barcodes. The average number of reads per non-reference barcode per sample was 5,655 and the median value was 3,442 (Figure S1). To avoid noise due to low counts entries with <= 20 reads were treated as missing data and removed prior to read summary reporting and downstream analyses.

#### Contamination rate

No instances of culture contamination were observed. Barcode cross-contamination rate was also measured and is defined as the total number of counts mapping to barcodes included in the full experimental design (library) but not expected to be present in that particular sample (given pair of forward/reverse Ion Torrent adapter Indices) [26]. The rate of barcode cross-contamination was tracked using samples seeded with a single pair of barcoded strains, such that the subsequent presence of other barcodes in these wells could be identified and quantified by sequencing. Barcode cross-contamination was low overall (0.269% +/-0.619%) and exhibited minor variation among sample sets (Day-0, 0.01% +/-0.007%; Day-50 salt, 0.030% +/-0.012%; Day-50 copper, 0.765% +/-0.990%).

#### Fitness calculations

The Malthusian fitness of focal barcoded line *i*, relative to the reference (d1H10), at experimental evolution generation *gn, m*_*i gn*_, was measured as,

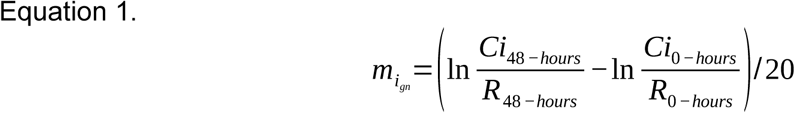

Where *Ci* and *R* refer to barcode counts for the focal barcode and reference barcode at fitness assay time 0-hours (initial mixtures) and time 48-hours (final overnight cultures), and 20 is the number of generations over 48 hours (two overnight cultures at 9.97 generations each – calculated from number of doublings based on optical density data) [27]. We use the standard equation *m=ln(w)* to convert Malthusian fitness values to Wrightian fitness values [28–30]. Hereafter, fitness, denoted by a *w* will refer to Wrightian fitness. The change in fitness of line *I* between Day-0 and Day-50, *Δw*_*i*_, was therefore computed as

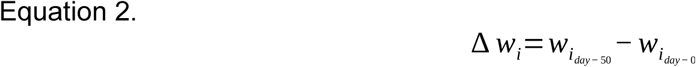

Where w_i day-0_ and w_i day-50_ are the line’s fitness relative to the ancestral reference at day-0 (before the experimental evolution) and at day-50 (end of experimental evolution) as measured from Equation 1. We assume no frequency dependent selection.

## Statistical analysis

### Analysis and visualization tools

R version 4.0.2 was used for all calculations, analyses, and figure generation [31]. Data processing uses base-R functionality supplemented with methods from the plyr package [32]. Statistical models with linear mixed effects utilize the lme4 [33] and lmerTest packages [34]. Power analyses were conducted using the pwr package [35]. Figures and tables were generated with ggplot2 [32] and sjPlot [36]; multi-panel figures were built using methods from grid [31], gridExtra [37], and cowPlot [38].

### Power analysis

Power analyses were conducted using root mean squared error (RMSE) and population standard deviation (PSD). RMSE for fitness change was 2.419 and PSD was 0.009. Consequently, we have 80% power to discern fitness differences of 0.306% between treatments and 80% power to identify fitness deviations from zero of 0.634% (Figure S2 B). We have 80% power to detect a fitness change (increase or decrease) of 2.163% for any individual barcode (Figure S2 A).

### Fitness change in 50 days of experimental evolution

Using the 80% power cutoff, we called individual barcoded yeast lines that increased (>= 2.163%) or decreased (<= 2.163%) in fitness over the 50-day experimental evolution. The effect of evolutionary treatment on fitness change was assessed using linear mixed-effects models with change in fitness as the response variable and treatment as the predictor variable. A random effect of line ID was placed on the model intercept (fitness change ∼ treatment + (1|line ID) + 0; family = gaussian). Separate models were run for each dataset by environment (salt lines in CM, salt lines in CM+NaCl, copper lines in CM, copper lines in CM+ CuSO_4_). This set of linear mixed effects models was repeated without the intercept term to assess fitness differences among treatments (rather than versus zero fitness change).

## Availability of data and materials

Data formatted for analysis and custom R scripts utilized for all data processing, statistical analyses, and figure generation are available from GitHub (https://github.com/VinceFasanello/FS_Code_Supplement). Instructions to reproduce the analyses and to confirm the results presented in this article are provided within. Yeast lines are available from the Fay Lab.

## Supporting information

Figure S1. Counts returned for sequenced libraries.

Figure S2. Power to detect fitness differences of different magnitudes as percent change from initial fitness.

Table S1. Lineages with fitness costs.

Table S2. Costs in the copper data.

Table S3. Costs in the salt data.

Table S4. Costs of generalization.

Table S5. Yeast strain barcodes

Table S6. Sequencing adapters and barcodes.

