## Supporting Figures for "Fluctuating selection facilitates the discovery of broadly effective but difficult to reach adaptive outcomes in yeast"

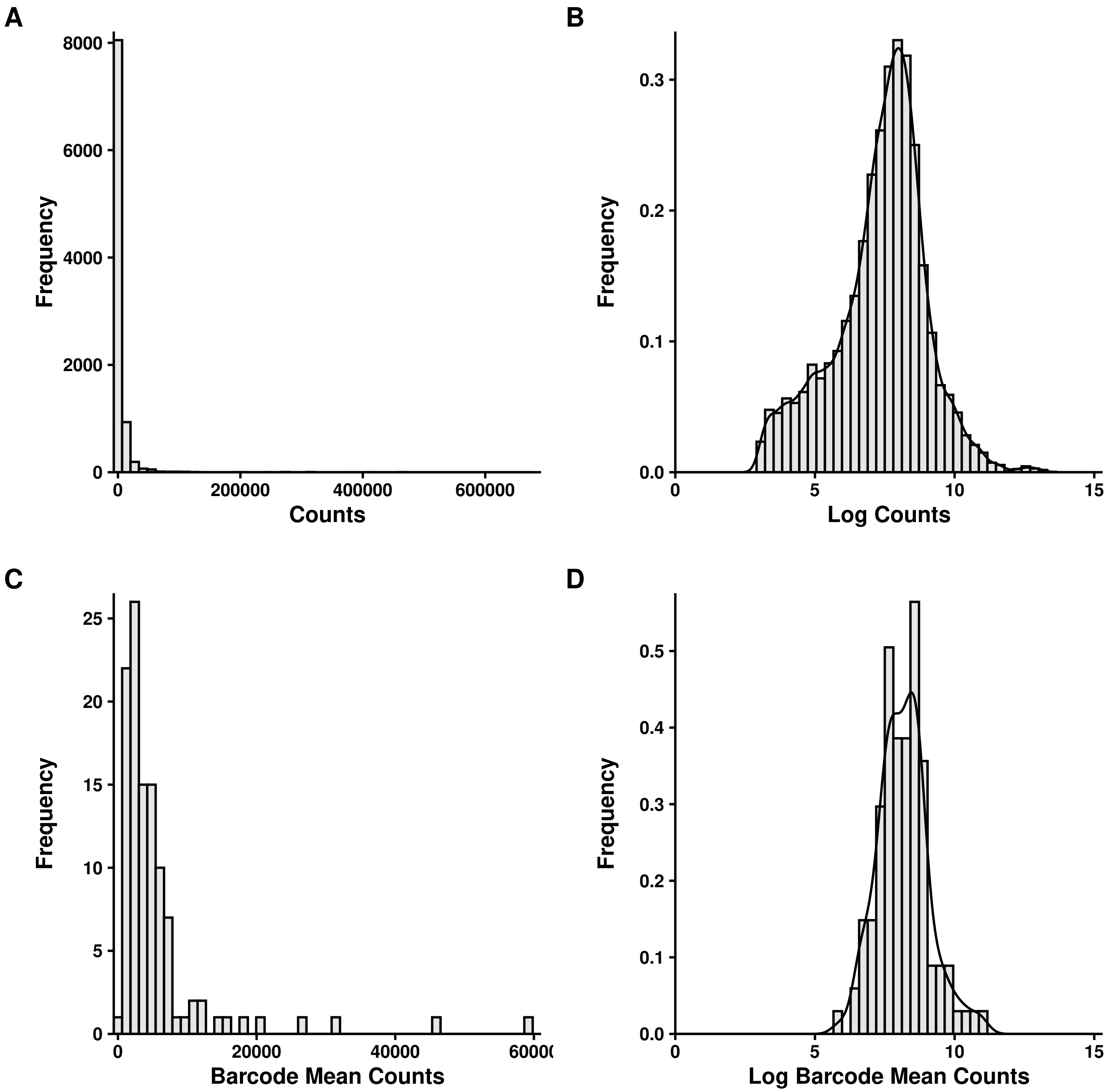


Figure S1: Counts returned for sequenced libraries. (A) Counts distribution for each barcoded lineage X sample (primer pair), omitting reference barcode data. (B) Log-counts distribution for each barcoded lineage X sample (primer pair), omitting reference barcode data. (C) Mean counts for each barcoded lineage across all samples, omitting reference barcode data. (D) Log-mean counts for each barcoded lineage across all samples, omitting reference barcode data.

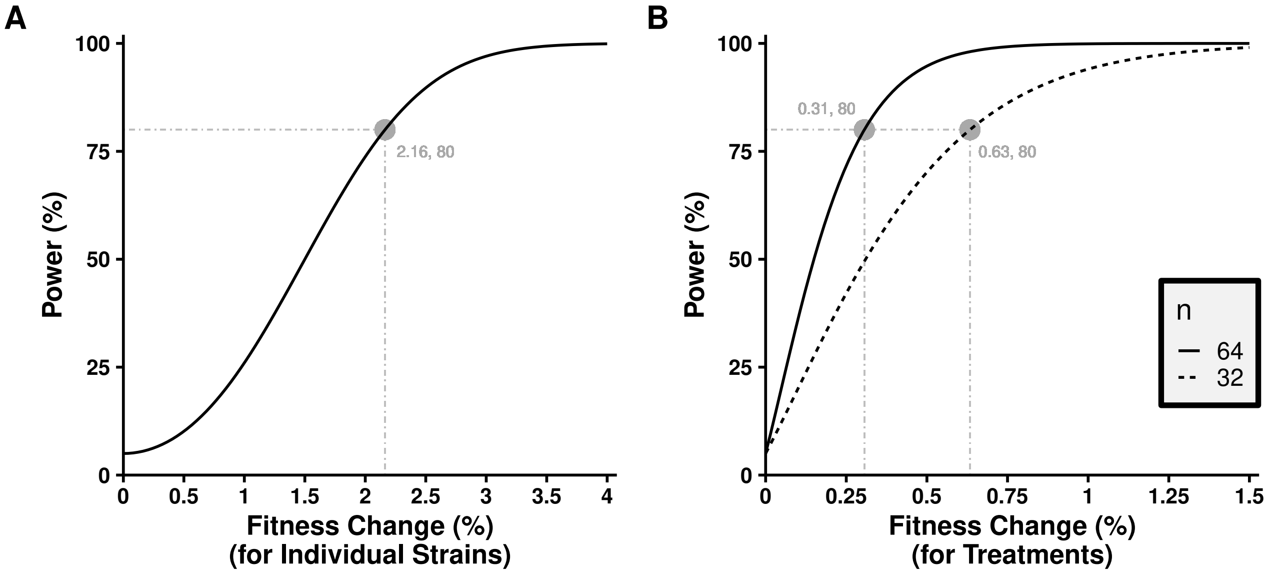


Figure S2: Power to detect fitness differences of different magnitudes as percent change from initial fitness. (A) power to detect fitness change for individual barcodes. (B) power to detect fitness change at the treatment level. N=64, solid line, is power to detect fitness difference between two treatments with 32 barcoded strains in each. N=32, dashed line, is power to detect fitness change for a single treatment with 32 barcoded strains.
